# Protein-glass microenvironments define electron transfer vulnerability in Leigh syndrome

**DOI:** 10.64898/2026.06.04.730076

**Authors:** Ji-Yong Sung, Jae-Ho Cheong

**Author notes:** Corresponding authors: **Ji-Yong Sung, Ph.D.**, Department of Neurosurgery, Seoul National University Bundang Hospital, Seoul National University College of Medicine, Seongnam-si, Republic of Korea **Jae-Ho Cheong, Ph.D., M.D.**, Department of Surgery, Yonsei University College of Medicine, Seoul 03722, Republic of Korea.

## Abstract

Leigh syndrome is the most common pediatric mitochondrial encephalopathy, yet the physical mechanisms linking diverse pathogenic mutations to respiratory-chain failure remain poorly understood. Here we show that Leigh syndrome mutations are not randomly distributed within human mitochondrial Complex I but are preferentially enriched near the electron-transfer axis connecting flavin mononucleotide and iron–sulfur cofactors. By integrating structural mutation mapping with residue-level free-volume analysis, packing-density measurements, and a protein glass index (PGI), we identify a distinct class of mutation-associated microenvironments characterized by reduced free volume, elevated packing density, and increased structural constraint. These protein-glass-like microenvironments were associated with elevated reorganization-energy proxies and increased transport vulnerability. Structure-informed Marcus analyses further identified a dominant bottleneck that was primarily associated with reduced electronic coupling arising from donor–acceptor separation, while local microenvironmental constraints further amplified transport sensitivity.

Structure-informed Marcus analyses identify the emergence of a dominant kinetic bottleneck within the iron–sulfur cluster network, and a Lindblad-based open quantum transport model indicates that local microenvironmental perturbations can propagate into network-level transport efficiency across the Complex I redox chain. Notably, pathogenic mutations preferentially accumulate in structural neighborhoods that are intrinsically sensitive to electron-transfer perturbation, suggesting that disease-associated variants may amplify pre-existing transport vulnerabilities embedded within the protein architecture. Collectively, our findings suggest a structural-energetic link between mutation landscapes, local protein-glass-like organization, and mitochondrial electron transport. We propose that Leigh syndrome can be viewed, in part, through a protein-glass lens in which pathogenic mutations preferentially map to structural-energy landscapes surrounding redox cofactors that are predicted to be vulnerableto electron-transfer perturbations. This framework provides a physical perspective for understanding genotype-to-phenotype convergence in mitochondrial disease and identifies local protein-glass analogous microenvironments as previously unrecognized determinants of respiratory-chain dysfunction.

## Introduction

Leigh syndrome is the most common pediatric mitochondrial encephalopathy and is characterized by progressive neurodegeneration, severe metabolic dysfunction, and early mortality.(*1*) Despite the identification of hundreds of pathogenic variants affecting mitochondrial oxidative phosphorylation, the mechanistic link between genotype and respiratory failure remains poorly understood.(*2*) Most studies have focused on biochemical defects, loss of catalytic activity, or impaired complex assembly.(*3*) However, these approaches do not fully explain why structurally diverse mutations distributed across multiple subunits can converge on a remarkably similar disease phenotype.(*4*) Mitochondrial Complex I occupies a central position in cellular bioenergetics, coupling electron transfer from NADH to ubiquinone with proton translocation across the inner mitochondrial membrane.(*5*) Electron transport within Complex I occurs through a chain of flavin and iron–sulfur redox centers separated by nanometer-scale distances. Efficient transfer requires a delicate balance among electronic coupling, reorganization energy, and environmental fluctuations.(*6–8*) Consequently, the local structural environment surrounding redox cofactors is expected to play a critical role in determining transport efficiency.(*9*) Yet, how disease-associated mutations alter this environment remains largely unknown.(*4*) Proteins are not static molecular scaffolds but dynamic energy landscapes that exhibit heterogeneous packing, local free-volume fluctuations, and glass-like structural constraints.(*10*) Increasing evidence suggests that biological function can be strongly influenced by these mesoscale structural properties. In densely packed regions, reduced free volume and restricted conformational fluctuations may alter local electrostatic environments, modify electron-transfer energetics, and perturb long-range transport pathways. (*8, 11*) Such effects are particularly relevant in respiratory complexes, where electron transfer occurs within structurally heterogeneous protein matrices. (*12*)

Here we propose that Leigh syndrome can be viewed, at least in part, as a disorder of protein-glass microenvironments surrounding the Complex I electron-transfer chain. (*12–14*) Rather than focusing exclusively on catalytic residues or protein stability, we investigate whether pathogenic mutations preferentially localize to structurally constrained regions that influence redox-center function.(*4*) To address this question, we developed a structure-based framework that integrates residue-level mutation mapping, local free-volume estimation, packing-density analysis, and a Protein Glass Index (PGI) derived from cryo-electron microscopy structures of human Complex I. Using this framework, we show that Leigh syndrome mutations are significantly enriched near the redox axis of Complex I and preferentially occupy densely packed structural environments characterized by reduced free volume and elevated packing density.(*15, 16*) These microenvironmental changes are associated with increased reorganization-energy proxies and the emergence of a pronounced electron-transfer bottleneck within the iron–sulfur cluster network.(*17, 18*) By combining Marcus-type electron-transfer analysis with open quantum transport modeling, we further demonstrate how local structural constraints can propagate into system-level transport defects. (*19–21*)

Our results support a mechanistic model in which pathogenic mutations reshape the glass-like structural landscape surrounding redox cofactors, thereby altering electron-transfer energetics and reducing transport efficiency. This perspective connects genetic variation, protein structural organization, and quantum transport within a unified framework, providing a new physical interpretation of mitochondrial disease and revealing protein-glass dynamics as a previously underappreciated determinant of respiratory-chain dysfunction. (*12, 18*)

## Results

### Leigh syndrome mutations are enriched near the electron-transfer axis of Complex I

To investigate whether Leigh syndrome mutations exhibit non-random spatial organization within mitochondrial Complex I, we mapped pathogenic variants onto the cryo-electron microscopy structure of the human enzyme (PDB 5XTD) and quantified their relationship to the major redox centers responsible for electron transport. A structural mutation atlas revealed that disease-associated variants were broadly distributed across multiple membrane and peripheral-arm subunits but showed a pronounced tendency to cluster around the electron-transfer pathway. (**Fig. 1A; Supplementary Table 1**) Rather than being uniformly dispersed throughout the complex, many mutations localized in regions surrounding flavin mononucleotide (FMN) and the iron–sulfur cluster chain connecting NADH oxidation to ubiquinone reduction. (**Supplementary Tables 1 and 2)**

**Figure 1.**
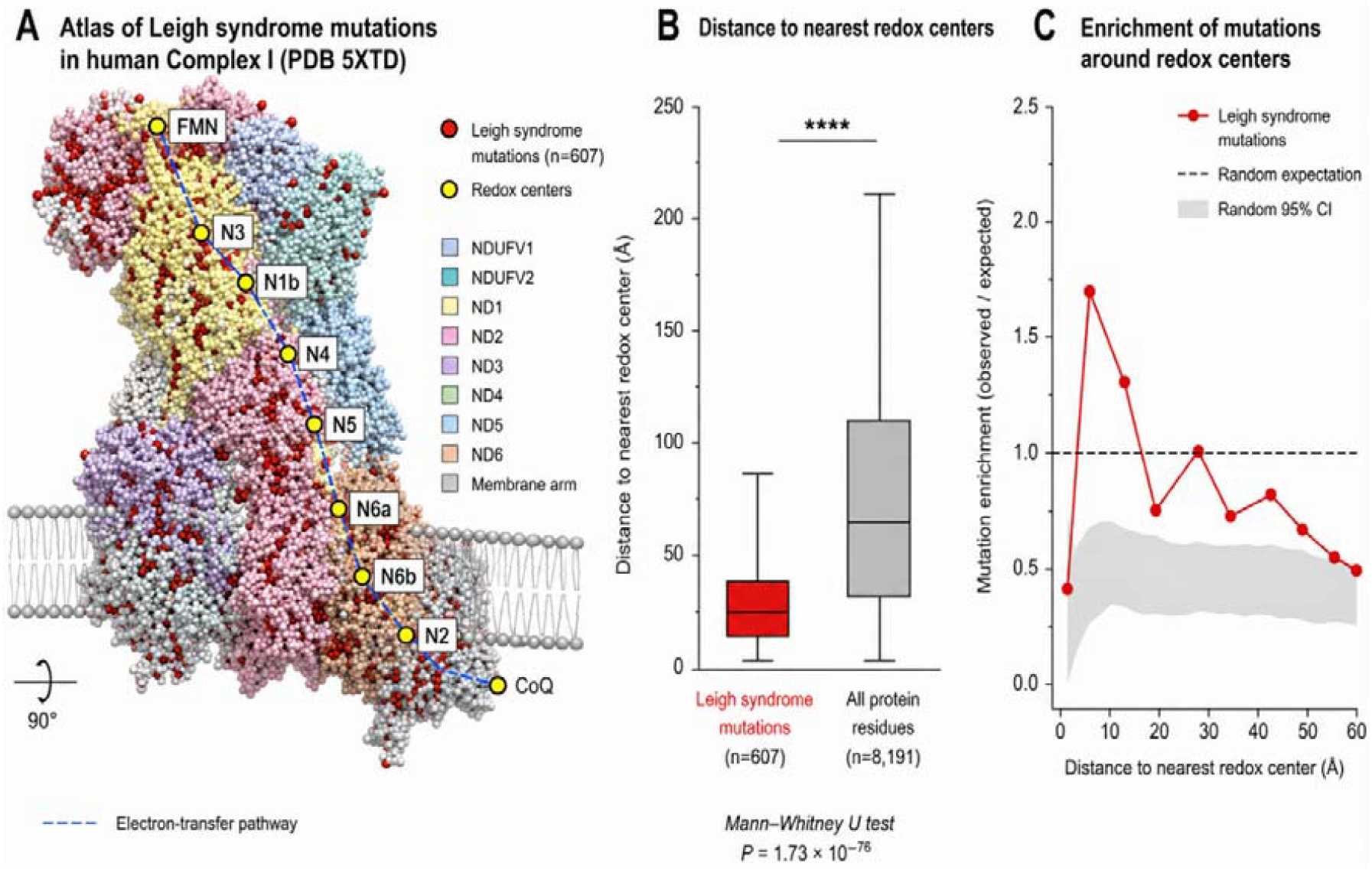
Leigh syndrome mutations preferentially localize near electron-transfer regions of human mitochondrial Complex I. (A) Structural atlas of Leigh syndrome-associated mutations mapped onto the human mitochondrial Complex I structure (PDB 5XTD). Red spheres indicate pathogenic Leigh syndrome mutations (n = 607), while yellow spheres denote electron-transfer redox centers, including FMN and Fe–S clusters (N3, N1b, N4, N5, N6a, N6b, and N2), as well as the terminal ubiquinone (CoQ) binding region. The blue dashed line illustrates the electron-transfer pathway from FMN to CoQ. Mutations are broadly distributed throughout Complex I but exhibit pronounced clustering near the electron-transfer axis. (B) Distribution of distances between Leigh syndrome mutations and the nearest redox center compared with all protein residues in Complex I. Leigh syndrome mutations are located significantly closer to redox-active sites than expected from the overall residue distribution (Leigh mutations, n = 607; all residues, n = 8,191). Box plots show median, interquartile range, and whiskers extending to 1.5× the interquartile range. Statistical significance was assessed using a two-sided Mann– Whitney U test (*P* = 1.73 × 10□□□). (C) Enrichment analysis of mutation frequency as a function of distance from the nearest redox center. Red circles indicate the observed enrichment of Leigh syndrome mutations relative to random expectation, whereas the gray shaded region represents the 95% confidence interval obtained from permutation-based random sampling. Mutations show the strongest enrichment within the structural neighborhood of redox centers (~5–15 Å), suggesting that pathogenic variants preferentially affect residues positioned near, but not directly on, the electron-transfer cofactors. Overall, these results demonstrate that Leigh syndrome mutations are non-randomly distributed within Complex I and preferentially target regions spatially associated with the electron-transfer machinery, supporting the hypothesis that perturbation of the local redox environment contributes to mitochondrial dysfunction in Leigh syndrome.

To quantify this spatial relationship, we calculated the distance between each mapped Leigh syndrome mutation and the nearest redox center (**Fig. 1B; Supplementary Table 3**). Mutation sites were significantly closer to the redox axis than expected from the distribution of all protein residues. The median distance of Leigh-associated mutation sites was approximately 30 Å, whereas the median distance across all residues exceeded 60 Å. This result indicates that pathogenic variants preferentially occupy structural neighborhoods surrounding electron-transfer cofactors rather than randomly sampling the available protein volume.

We next examined mutation enrichment as a function of distance from the redox chain. Leigh syndrome variants exhibited a marked enrichment within the immediate vicinity of redox centers, with mutation density progressively decreasing at larger distances (**Fig. 1C; Supplementary Table 3**). The strongest enrichment was observed within the first structural shell surrounding FMN and the iron–sulfur cluster chain, indicating that regions directly associated with electron transfer constitute preferential targets of pathogenic variation. Representative disease-associated variants further support this non-random spatial organization (**Supplementary Table 1**). Notably, ND6 Y42C and multiple pathogenic variants affecting the NDUFS1 subunit were located within structurally constrained microenvironments adjacent to major redox cofactors (**Supplementary Tables 1 and 2**). Although these mutations occur in distinct subunits and are associated with diverse molecular defects, their preferential localization near the electron-transfer axis suggests a convergent pathogenic mechanism. Specifically, disruption of the densely packed protein environment surrounding redox-active centers may alter local energetic landscapes, perturb electron-transfer efficiency, and ultimately compromise long-range charge transport through Complex I.

Consistently, these findings demonstrate that Leigh syndrome mutations are spatially biased toward the electron-transfer architecture of Complex I. This enrichment cannot be explained solely by subunit composition or residue abundance and instead suggests that disease-associated variants preferentially affect regions that are functionally coupled to electron transport. The observed spatial organization provides the structural foundation for subsequent analyses examining how mutation-associated microenvironmental changes influence electron-transfer energetics and transport dynamics.

### Leigh syndrome mutations occupy densely constrained protein microenvironments surrounding the Complex I redox axis

Having established that Leigh syndrome mutations are preferentially localized near the electron-transfer pathway of Complex I (**Fig. 1**), we next investigated whether these mutation-associated regions exhibit distinctive structural microenvironmental properties. To address this question, we quantified local free volume, packing density, and a Protein Glass Index (PGI) for residues located within the structural neighborhood of major redox centers. Analysis of local free-volume distributions revealed substantial heterogeneity along the redox axis (**Fig. 2A**). Regions surrounding FMN and proximal iron–sulfur clusters exhibited relatively large accessible free-volume estimates, whereas downstream portions of the electron-transfer chain displayed progressively reduced free-volume signatures. This pattern was accompanied by a reciprocal increase in local packing density, indicating the presence of structurally constrained environments surrounding specific redox centers. To integrate these structural descriptors, we calculated a Protein Glass Index (PGI), which captures the balance between local free volume, packing density, and residue-level spatial organization. Mapping PGI values onto the redox pathway demonstrated a gradual transition from relatively flexible environments near the NADH oxidation site toward increasingly constrained environments within the distal iron–sulfur cluster network (**Fig. 2B**). Notably, several redox centers located in mutation-enriched regions exhibited among the lowest PGI values observed across the electron-transfer chain. Under the PGI definition used here, lower PGI values correspond to densely packed and structurally constrained microenvironments characterized by reduced free volume and increased packing density.

**Figure 2.**
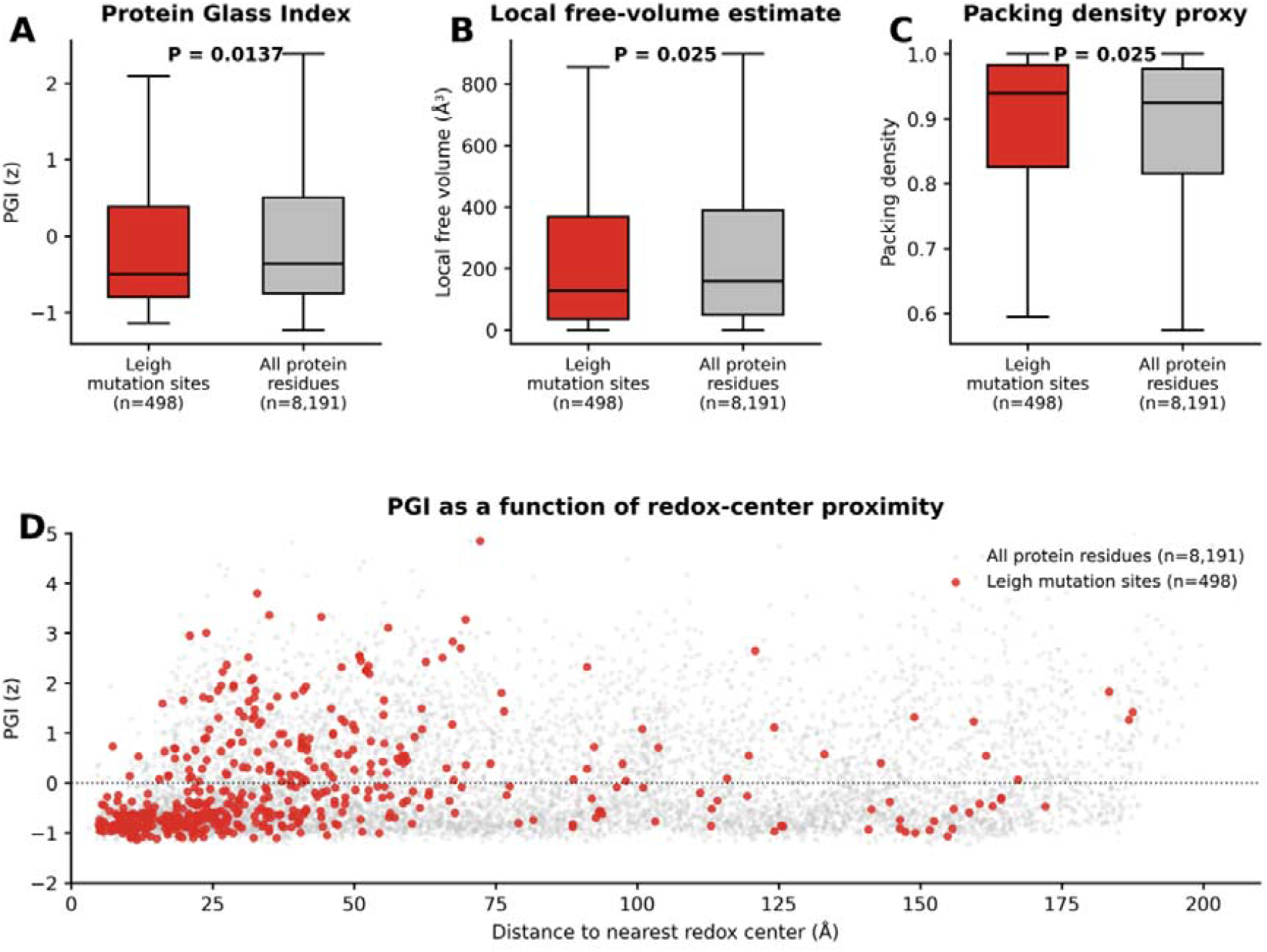
Leigh syndrome mutation sites occupy densely packed and structurally constrained microenvironments in human mitochondrial Complex I. (A) Protein Glass Index (PGI) values were compared between Leigh syndrome mutation sites and all protein residues in human Complex I. Leigh syndrome mutation sites exhibited significantly lower PGI values than the overall residue population, indicating preferential localization within densely packed and structurally constrained microenvironments. (B) Local free-volume estimates were lower at Leigh syndrome mutation sites compared with all protein residues, indicating reduced local structural space around pathogenic mutation positions. (C) Packing-density proxy values were higher at Leigh syndrome mutation sites, suggesting that pathogenic variants preferentially reside in densely packed structural environments. (D) PGI values plotted as a function of distance to the nearest redox center. The x-axis was verified using the full distance range of the dataset (0–210 Å). No regression line was applied. Boxplots show the median and interquartile range; whiskers indicate 1.5×IQR. P values were calculated using two-sided Mann–Whitney U tests. PGI was calculated as

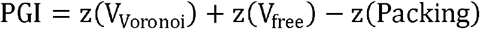

where V_Voronoi_ denotes the residue-level Voronoi volume estimate obtained from Monte Carlo Voronoi tessellation, V_free_ denotes the local free-volume estimate, and Packing denotes the local packing-density proxy. Under this definition, higher PGI values indicate structurally permissive microenvironments characterized by larger free volume and lower packing density, whereas lower PGI values indicate densely packed and structurally constrained regions with reduced configurational freedom.

Direct comparison of mutation-associated neighborhoods and background structural regions further demonstrated that Leigh syndrome mutations preferentially occur within densely packed environments characterized by reduced free volume and elevated packing density (**Fig. 2C**). These observations suggest that pathogenic variants are not randomly distributed across the structural landscape of Complex I but instead preferentially target regions that already possess limited conformational freedom. To evaluate the combined effect of these structural features, we derived a dense-constraint score representing the relative degree of local microenvironmental restriction. Redox centers exhibiting elevated dense-constraint scores were strongly associated with mutation-enriched regions and displayed reduced PGI values (**Fig. 2D**). This relationship indicates that disease-associated variants accumulate within low-PGI structural environments that are intrinsically less permissive to local fluctuations.

Collectively, these results reveal that Leigh syndrome mutations preferentially occupy protein-glass microenvironments characterized by reduced free volume, increased packing density, and elevated structural constraint. Importantly, these effects are concentrated around the electron-transfer axis of Complex I, suggesting that pathogenic mutations may influence respiratory-chain function not only through direct biochemical perturbations but also through alterations of the local structural-energy landscape. These findings provide a mechanistic basis for examining how mutation-associated microenvironmental constraints affect electron-transfer energetics and transport dynamics in subsequent analyses.

### Protein-glass microenvironments generate a localized electron-transfer bottleneck within Complex I

To determine whether mutation-associated structural constraints influence electron-transfer energetics, we next examined the local microenvironment surrounding each major redox center of Complex I and evaluated its impact on electron-transfer parameters. Structural descriptors derived from local free volume, packing density, and Protein Glass Index (PGI) were integrated to characterize the microenvironmental landscape along the electron-transfer axis. A redox-center–resolved analysis revealed substantial heterogeneity in structural microenvironmental properties across the electron-transfer chain (**Fig. 3A**). FMN and proximal iron–sulfur clusters exhibited relatively elevated PGI and free-volume signatures, whereas downstream redox centers were associated with progressively denser structural environments characterized by reduced free volume and increased packing density. These observations indicate that the electron-transfer pathway traverses regions with markedly different local structural constraints rather than a uniform protein matrix. We next estimated hop-specific reorganization-energy proxies (λ) for sequential electron-transfer steps connecting the major redox centers (**Fig. 3B**). λ proxy values increased along the transport pathway and reached their highest levels within the distal portion of the iron–sulfur cluster network. This trend suggests that structurally constrained microenvironments may impose larger energetic penalties on electron transfer by increasing the degree of environmental reorganization required during charge movement.

**Figure 3.**
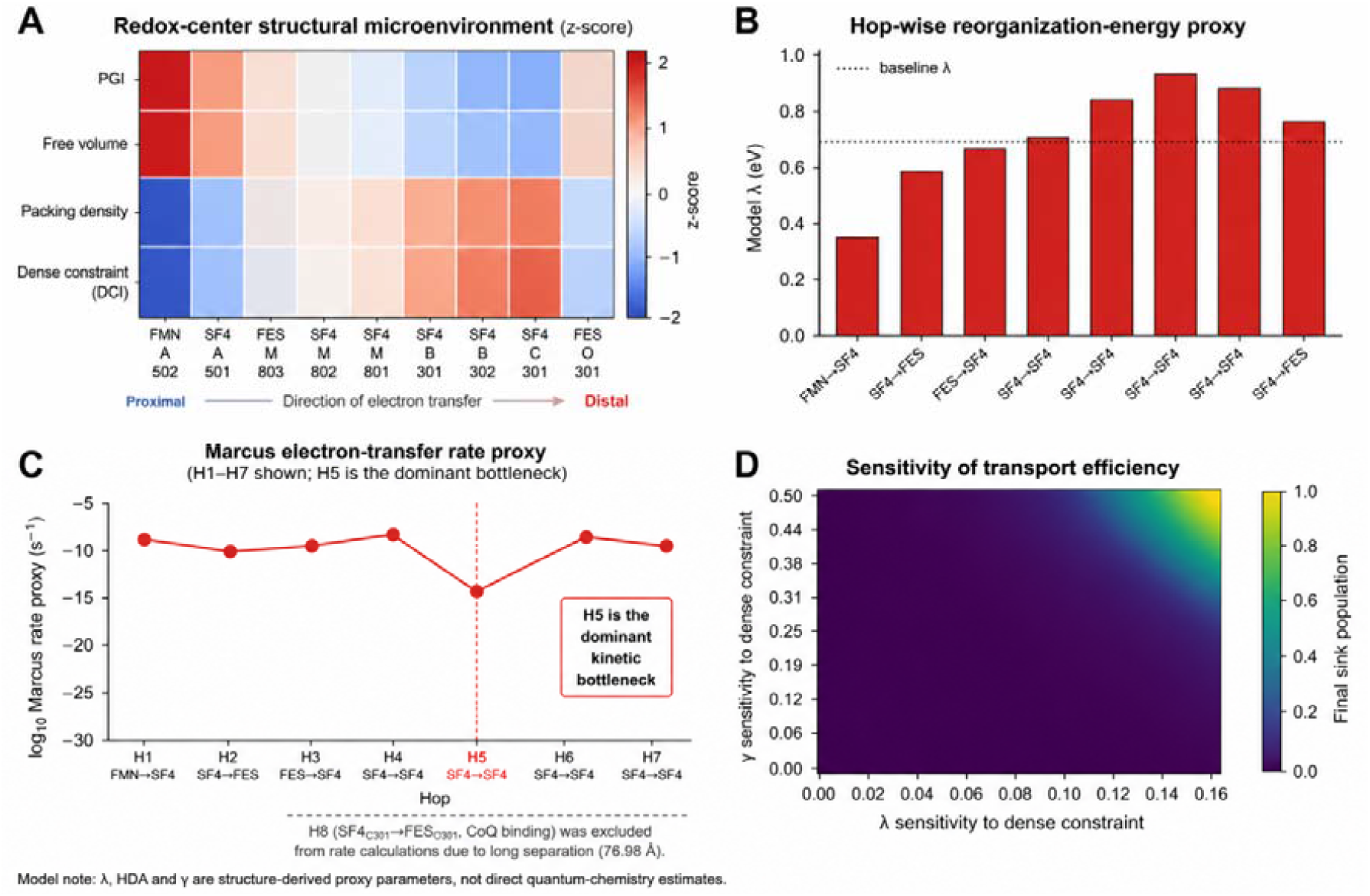
Structural microenvironment, electron-transfer energetics, and transport sensitivity along the Complex I redox chain. (A) Z-score-normalized structural microenvironment properties surrounding the major redox centers of human mitochondrial Complex I (PDB 5XTD). Protein Glass Index (PGI), local free-volume estimate, packing-density estimate, and Dense Constraint Index (DCI) were calculated for residues located within 15 Å of each redox center and standardized across all redox sites. PGI and free-volume signatures progressively decreased along the electron-transfer pathway, whereas packing density and DCI increased toward distal Fe–S clusters. Because PGI represents structural permissiveness, higher PGI values correspond to microenvironments with greater free volume and lower packing density, whereas higher DCI values indicate increased structural constraint characterized by reduced free volume, elevated packing density, and lower structural permissiveness. Thus, PGI and DCI exhibit an inverse relationship across the Complex I redox chain. (B) Hop-wise reorganization-energy (λ) proxy along the electron-transfer pathway. λ proxy values were derived from local structural descriptors surrounding adjacent redox centers and represent relative estimates of environmental reorganization costs associated with electron transfer. Reorganization-energy proxies increased along the redox pathway and reached their highest values within the distal portion of the Fe–S cluster network, indicating progressively constrained transport environments. (C) Marcus electron-transfer rate proxy for structurally adjacent redox hops. Relative electron-transfer rates were estimated using structure-derived proxy parameters for electronic coupling and reorganization energy. Only biologically plausible nearest-neighbor hops (H1– H7) are shown. The H5 transition (SF4_M_801 → SF4_B_301) exhibited the lowest predicted transfer rate and was identified as the dominant kinetic bottleneck within the analyzed redox chain. The terminal long-distance transition SF4_C_301 → FES_O_301 (76.98 Å) was excluded from rate calculations because its separation substantially exceeded the distance range typically associated with efficient biological electron transfer. (D) Sensitivity landscape of transport efficiency. Final sink population obtained from Lindblad-based open quantum transport simulations is shown as a function of λ sensitivity and γ sensitivity to local dense constraint. Warmer colors indicate higher transport efficiency, whereas cooler colors indicate reduced transport efficiency. The landscape illustrates the combined influence of environmental reorganization and dephasing sensitivity on network-level electron transport and identifies parameter regimes associated with reduced transport performance. Model note: λ, HDA, and γ represent structure-derived proxy parameters inferred from local protein microenvironment properties and should not be interpreted as experimentally calibrated quantum-mechanical quantities.

To evaluate the functional consequences of these changes, we calculated Marcus electron-transfer rate proxies for structurally adjacent redox hops (**Fig. 3C**). Most transfer steps exhibited comparable rate estimates; however, a pronounced decrease was observed for the transition between the SF4_M_801 and SF4_B_301 redox centers (H5). This hop displayed the lowest predicted transfer rate in the entire network (log10 rate proxy ≈ −13.85), identifying it as the dominant kinetic bottleneck within the Complex I electron-transfer chain.

Importantly, analysis of the underlying Marcus parameters indicated that the dominant contribution arose from reduced electronic coupling associated with the relatively long donor–acceptor separation. Elevated reorganization-energy proxy values contributed secondarily to the transport limitation. Thus, the H5 bottleneck is primarily consistent with distance-dependent electron-transfer theory, while local microenvironmental constraints further amplify the severity of the transport defect.

To evaluate how local structural constraints influence network-level transport behavior, we next examined the sensitivity of electron-transport efficiency to microenvironment-derived perturbations using Lindblad-based open quantum transport simulations (**Fig. 3D**). Transport efficiency was quantified as the final sink population and evaluated across a range of λ and γ sensitivities to local dense constraint. The resulting sensitivity landscape revealed that transport efficiency remained relatively stable across low-sensitivity regimes but declined substantially as both reorganization-energy sensitivity and dephasing sensitivity increased simultaneously. These findings indicate that structurally constrained microenvironments can amplify transport vulnerability through coupled energetic and dynamical effects.

Taken together, these findings provide a mechanistic link between structurally constrained protein-glass microenvironments, electron-transfer energetics, and network-level transport vulnerability. The results indicate that local protein-glass microenvironments are not passive structural features but active determinants of electron-transfer energetics. By increasing local energetic constraints, elevating reorganization-energy penalties, and enhancing transport sensitivity to environmental perturbation, these microenvironments create localized vulnerabilities within the Complex I redox chain that may amplify the functional consequences of Leigh syndrome mutations.

Notably, the dominant bottleneck identified here was confined to a specific electron-transfer step rather than being uniformly distributed across the entire redox network. This observation suggests that Leigh syndrome may arise from the selective amplification of pre-existing transport vulnerabilities embedded within the structural-energy landscape of Complex I, providing a direct physical connection between protein-glass organization and mitochondrial electron-transfer dysfunction.

### Structural characterization and rescue potential of the dominant electron-transfer bottleneck

Having established the H5 transition, connecting SF4_M_801 and SF4_B_301, as the principal rate-limiting step within the Complex I electron-transfer network (**Fig. 3C**), we next investigated the structural origin, robustness, and theoretical reversibility of this transport bottleneck. The experimentally resolved redox-center network revealed that H5 occupies a distinctive position within the Fe–S cluster chain, where relatively long donor–acceptor separation coincides with elevated structure-derived reorganization-energy proxies (**Fig. 4A**). In contrast to neighboring electron-transfer steps, the combination of these unfavorable structural parameter’s places H5 in a region of heightened transport sensitivity. Because the terminal electron-transfer step linking SF4_C_301 and FES_O_301 exhibited an exceptionally large donor–acceptor separation (76.98 Å), it was deemed structurally incompatible with direct electron transfer and was therefore excluded from subsequent kinetic analyses. To determine the relative contribution of each redox hop to overall transport limitation, we ranked bottleneck scores across all structurally adjacent electron-transfer steps (**Fig. 4B**). Following exclusion of the long-distance artifact hop, H5 remained the largest bottleneck in the network, exceeding all neighboring transitions. This result confirms that the kinetic vulnerability identified in Fig. 3 is not a consequence of global transport slowing but instead reflects a localized constraint embedded within a specific segment of the Fe–S cluster pathway. We next assessed the robustness of this bottleneck by systematically perturbing the two principal determinants of Marcus electron transfer: electronic coupling and reorganization energy. Across a broad parameter space, H5 consistently remained within a low-efficiency transport regime (**Fig. 4C**), indicating that its bottleneck behavior is structurally stable rather than dependent on a narrow set of model assumptions. Notably, predicted transfer efficiency was substantially more sensitive to increases in electronic coupling than to equivalent changes in reorganization energy, suggesting that donor–acceptor communication represents the dominant limiting factor at this step. Consistent with classical Marcus theory, transport efficiency was predominantly governed by electronic coupling. Reorganization-energy changes modulated the severity of the bottleneck but did not determine its location within the redox chain.

**Figure 4.**
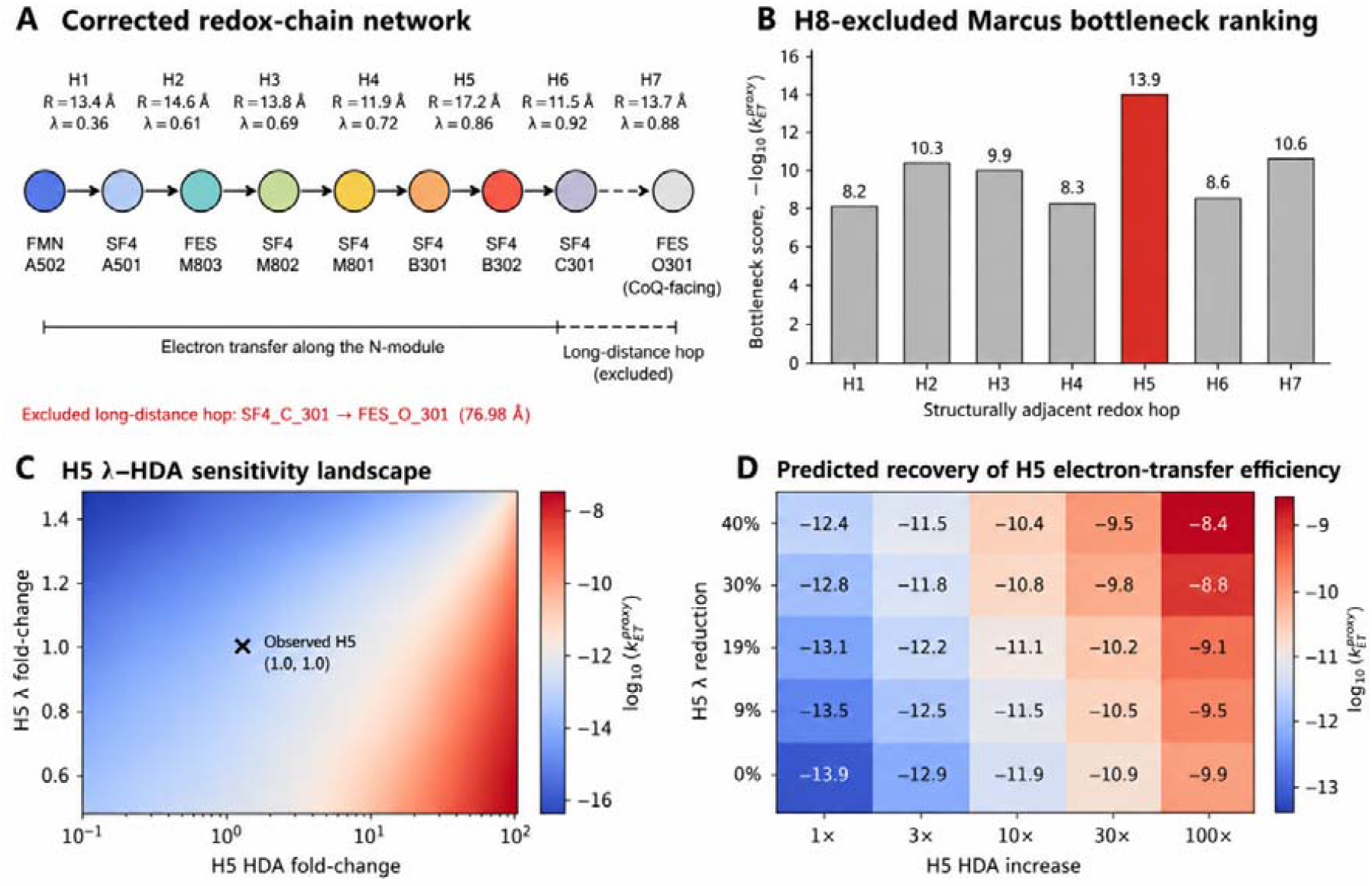
Structural identification and rescue analysis of the dominant electron-transfer bottleneck in human mitochondrial Complex I. (A) Corrected redox-chain network derived from the cryo-EM structure of human mitochondrial Complex I (PDB ID: 5XTD). Structurally adjacent redox centers were connected according to the experimentally resolved electron-transfer pathway. Donor– acceptor distances R_DA_ and structure-derived reorganization-energy proxies ((\lambda)) are shown for each hop. The non-physical long-distance transition SF4_C_301 → FES_O_301 (76.98 Å) was excluded from kinetic analyses because its separation exceeds the distance range typically associated with biologically relevant electron transfer. (B) Marcus bottleneck ranking for structurally adjacent redox hops following exclusion of the long-distance artifact hop. Bottleneck scores were calculated as 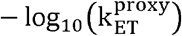, where larger values indicate slower predicted electron transfer. The H5 transition (SF4_M_801 → SF4_B_301) exhibited the highest bottleneck score, identifying it as the dominant kinetic constraint within the analyzed redox chain. (C) Sensitivity landscape of the H5 bottleneck. The structure-derived Marcus electron-transfer rate proxy was recalculated across a broad range of electronic-coupling H_DA_ and reorganization-energy λ perturbations. Warmer colors indicate higher predicted electron-transfer efficiency. The observed H5 parameter set is indicated by the black cross. The landscape demonstrates that H5 remains kinetically unfavorable across a wide parameter space and is particularly sensitive to increases in electronic coupling. (D) Predicted recovery of H5 electron-transfer efficiency. The Marcus rate proxy was recalculated following systematic increases in electronic coupling H_DA_ and reductions in reorganization energy λ. Values represent 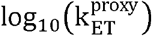. Progressive enhancement of electronic coupling and reduction of reorganization energy substantially improved the predicted transfer efficiency, suggesting that the H5 bottleneck is theoretically reversible through local structural or energetic optimization. All rate values represent relative Marcus-inspired electron-transfer proxies derived from structural parameters and should not be interpreted as absolute experimental electron-transfer rates.

Finally, rescue simulations were performed to evaluate whether the H5 bottleneck could be theoretically alleviated through local microenvironmental modification (**Fig. 4D**). Simultaneous enhancement of electronic coupling and reduction of reorganization-energy penalties progressively improved the predicted electron-transfer rate. The largest perturbations produced improvements exceeding several orders of magnitude in the Marcus rate proxy, demonstrating that the bottleneck is not an irreversible property of the redox architecture itself but rather emerges from the local structural-energy landscape surrounding the H5 region. Together, these findings extend the observations of Fig. 3 by showing that the dominant electron-transfer bottleneck in Complex I is structurally localized, kinetically robust, and theoretically reversible. The results support a model in which mutation-associated protein-glass microenvironments do not merely alter local structural properties but can selectively amplify pre-existing transport vulnerabilities embedded within the Complex I electron-transfer network.

## Discussion

The prevailing view of Leigh syndrome has largely focused on defects in oxidative phosphorylation, impaired enzyme activity, and disruption of respiratory-complex assembly. (*22*) Although these mechanisms undoubtedly contribute to disease pathogenesis, they do not fully explain how a diverse collection of mutations distributed across multiple Complex I subunits can converge on a common bioenergetic phenotype.(*23*) Our results suggest an additional layer of organization: the local structural microenvironment surrounding the electron-transfer pathway.(*24*) By integrating mutation mapping, structural microenvironment analysis, Marcus-type electron-transfer modeling, and open quantum transport simulations, we found that Leigh syndrome mutations are not randomly distributed throughout Complex I.

Instead, pathogenic variants preferentially localize near the redox axis and occupy densely packed structural environments characterized by reduced free volume, increased packing density, and lower Protein Glass Index (PGI) values. Under the PGI formulation used in this study, lower PGI values correspond to densely packed and structurally constrained microenvironments with reduced configurational freedom, whereas higher PGI values indicate more permissive local environments characterized by greater free volume and lower packing density.

These observations suggest that disease-associated mutations are enriched within regions that are particularly sensitive to fluctuations in electron-transfer energetics. A central finding of this study is that local structural constraints can be translated into altered electron-transfer parameters. Within the framework of Marcus theory, electron-transfer efficiency depends critically on electronic coupling and reorganization energy. The structural microenvironments identified here were associated with increased reorganization-energy proxies and reduced transfer-rate estimates, indicating that densely packed regions may impose energetic penalties on electron movement through the iron–sulfur cluster network.(*25*)

Notably, the dominant H5 bottleneck was primarily associated with reduced electronic coupling arising from donor–acceptor separation rather than the largest reorganization-energy proxy. This is the expected behavior under classical Marcus and Moser–Dutton electron-transfer theory, in which electronic coupling decays steeply with distance. We therefor interpret that protein-glass-like microenvironments act mainly as amplifiers of an existing transport vulnerability.The transport-sensitivity analysis further suggests that these structural constraints influence not only local electron-transfer rates but also the robustness of network-level transport behavior. Increased sensitivity of sink population to λ- and γ-dependent perturbations indicates that densely constrained protein-glass microenvironments may amplify the impact of otherwise modest environmental fluctuations. This observation supports the concept that local structural organization can propagate into system-level transport vulnerability within the Complex I redox network. Importantly, these effects were not uniformly distributed across the redox chain. Instead, the analysis identified a dominant bottleneck associated with a specific redox hop, suggesting that pathogenic mutations may amplify pre-existing vulnerabilities within the electron-transfer architecture of Complex I.(*7, 26*)

This observation has important conceptual implications. Classical descriptions of mitochondrial disease often assume that pathogenic mutations reduce respiratory-chain performance in a relatively uniform manner. Our results support an alternative picture in which disease emerges from localized perturbations within a structurally heterogeneous transport network. In this view, the functional consequences of a mutation depend not only on its biochemical identity but also on its position within the structural-energy landscape of the protein. Mutations occurring within densely constrained microenvironments may exert disproportionately large effects because they alter regions that already operate near critical transport limits. (*27*)

The protein-glass perspective provides a useful framework for understanding these phenomena. Proteins are increasingly recognized as heterogeneous materials that exhibit glass-like properties, including restricted conformational dynamics, local packing heterogeneity, and non-uniform energy landscapes.(*28, 29*) Although such concepts have been widely explored in enzymology and biophysics, their relevance to mitochondrial disease has received comparatively little attention. Our findings suggest that local glass-like structural constraints may influence electron-transfer behavior by modulating the balance between environmental fluctuations and transport energetics.(*30*) From this perspective, Leigh syndrome may be viewed not solely as a disorder of protein composition but also as a disorder of protein-glass organization.(*31*) The integration of Marcus-type analysis with open quantum transport modeling further extends this interpretation. While the present transport simulations should be regarded as structure-informed proxy models rather than ab initio quantum-dynamical calculations, they provide a mechanistic bridge linking local structural constraints to system-level transport behavior. The emergence of reduced transport efficiency in mutation-constrained models suggests that microenvironmental perturbations can propagate across the redox network and influence global electron-flow dynamics. This framework naturally connects molecular pathology with concepts from quantum transport theory and open-system dynamics.(*32, 33*)

Several limitations should be considered. First, the free-volume, packing-density, and PGI measurements represent structure-derived proxy quantities inferred from static cryo-electron microscopy coordinates. Additional validation using molecular-dynamics ensembles, cavity-volume calculations, or experimental measurements of conformational fluctuations would strengthen the physical interpretation of these parameters. Second, the Marcus and transport analyses employ simplified proxy representations of electronic coupling and environmental reorganization. Future studies incorporating atomistic electronic-structure calculations, QM/MM approaches, or experimentally constrained redox parameters may refine the quantitative estimates reported here. Finally, although our analyses focus on human Complex I, the general framework may be applicable to other respiratory-chain complexes and mitochondrial disorders.

Despite these limitations, the present study establishes a conceptual link between pathogenic mutation landscapes, protein-glass microenvironments, and electron-transfer dynamics in mitochondrial Complex I. Rather than viewing Leigh syndrome exclusively as a consequence of defective oxidative-phosphorylation components, our results support a model in which disease-associated mutations reshape the structural-energy landscape surrounding the redox chain, thereby altering electron-transfer energetics and transport efficiency. This perspective provides a unifying physical framework that connects genotype, protein architecture, and bioenergetic dysfunction, and suggests that modulation of local structural constraints may represent an unexplored avenue for therapeutic intervention in mitochondrial disease.

## Methods

### Structural framework of the mitochondrial Complex I electron-transfer chain

Human mitochondrial respiratory Complex I was analyzed using the cryo-electron microscopy structure deposited in the Protein Data Bank (PDB ID: 5XTD).(*34*) This structure was selected because it provides near-complete atomic coordinates for the N-module electron-transfer pathway, including flavin mononucleotide (FMN) and all major iron–sulfur (Fe–S) clusters involved in long-range electron transport. The redox centers included in the analysis were FMN_A_502, SF4_A_501, FES_M_803, SF4_M_802, SF4_M_801, SF4_B_301, SF4_B_302, SF4_C_301, and FES_O_301. These cofactors constitute the principal electron-transfer chain connecting the NADH oxidation site to the ubiquinone reduction region. (*35*) For each redox center, three-dimensional centroid coordinates were calculated from all atoms belonging to the corresponding cofactor. Pairwise donor–acceptor distances R_DA_were calculated using Euclidean geometry:

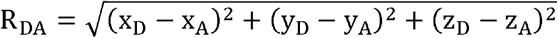

where (D) and (A) denote donor and acceptor redox centers, respectively. The resulting redox-chain topology served as the structural scaffold for all subsequent analyses.

### Mapping of Leigh syndrome mutations onto the Complex I structure

Pathogenic variants associated with Leigh syndrome were collected from curated mitochondrial disease databases and published literature.(*36*) Mutations were mapped onto the 5XTD structure through residue-number correspondence. Residues lacking resolved coordinates were excluded. For each residue i, the shortest distance to any redox center was calculated:

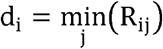

where R_ij_ denotes the distance between residue i and redox center j. Mutation enrichment near the electron-transfer chain was evaluated by comparing the observed mutation distribution with the distribution expected from all structurally resolved residues.(*37*)

### Quantification of local glass-like structural disorder

To characterize the heterogeneous structural environment surrounding redox centers, we introduced a Protein Glass Index (PGI). The conceptual basis of PGI derives from glass physics, where local packing heterogeneity and configurational restriction generate rugged energy landscapes containing multiple metastable states. We hypothesized that analogous structural heterogeneity may influence electron-transfer energetics within mitochondrial Complex I. For each residue, local structural descriptors were calculated within a sphere of radius 15 Å and independently standardized as z-scores:

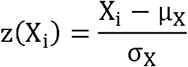

where X_i_ denotes the local structural descriptor and μ_X_ and σ_X_ represent the global mean and standard deviation across all residues. PGI was calculated by integrating standardized descriptors reflecting local structural permissiveness, including Voronoi volume, free volume, and packing density:

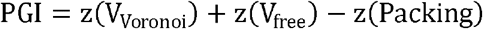

Under this definition, higher PGI values indicate structurally permissive microenvironments characterized by larger free volume, lower packing density, and greater configurational flexibility. Conversely, lower PGI values indicate densely packed and structurally constrained regions with reduced configurational freedom. Thus, PGI should be interpreted as a disorder/permissiveness index rather than a direct measure of structural constraint. For redox-center analyses, PGI values were averaged across all residues located within 15 Å of each cofactor centroid.

### Estimation of local free volume

Local free volume was estimated as a proxy for structural flexibility. For each residue-centered neighborhood, occupied volume was calculated from atomic van der Waals radii. Local free volume was then defined as (*39*)

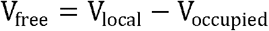

where V_local_ denotes the total neighborhood volume and V_occupied_ denotes the cumulative atomic volume. Although this approximation does not explicitly model solvent-accessible cavities, it provides a consistent relative measure of local configurational freedom across the protein structure. All free-volume measurements were converted to z-scores before inter-center comparison.

### Packing-density analysis

Packing density was used to quantify local structural compactness.(*40*) For each neighborhood,

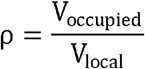

was calculated. Higher values correspond to denser packing and reduced local flexibility. Because PGI, free volume, and packing density possess different physical units, all values were standardized independently before integration or visualization.

### Dense-constraint index

To summarize multiple structural properties within a single metric, we defined a Dense Constraint Index (DCI). The index integrates local structural permissiveness, available free volume, and packing density:

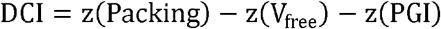

where Packing denotes the local packing-density estimate, V_free_ denotes the local free-volume estimate, and PGI denotes the Protein Glass Index. Because PGI is interpreted as a structural permissiveness index in this study, lower PGI values contribute positively to DCI. Higher DCI values therefore indicate densely packed and structurally constrained environments characterized by increased packing density, reduced free volume, and reduced structural permissiveness. Lower DCI values indicate comparatively flexible environments. The DCI was used to identify structurally restricted regions along the electron-transfer axis.

### Structure-derived reorganization-energy proxy

Classical Marcus theory predicts that electron-transfer rates depend strongly on the reorganization energy λ, which describes the energetic cost of environmental rearrangement during electron transfer.(*41*) Because direct quantum-chemical estimation of λ for the full Complex I structure remains computationally challenging, we derived a structure-based proxy. For each redox hop, the reorganization-energy proxy was modeled as a function of local structural constraints:

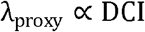

Because higher DCI values correspond to densely packed and structurally constrained microenvironments, the reorganization-energy proxy was modeled as increasing with increasing DCI. Thus, larger λ values reflect greater local structural reorganization costs associated with constrained protein environments. Consequently, densely packed and structurally constrained microenvironments are predicted to exhibit elevated reorganization-energy penalties.

Consequently, densely packed and structurally constrained microenvironments are predicted to exhibit elevated reorganization-energy penalties. (*42*)

### Electronic-coupling proxy

Electronic coupling was modeled using an exponential distance-decay approximation:

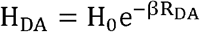

where (H_0_) is a scaling constant, R_DA_ is the donor–acceptor distance, and (\beta) is the distance-decay coefficient. This approximation reflects the established decrease in electronic overlap with increasing donor–acceptor separation. The resulting values are interpreted as relative electronic-coupling proxies rather than direct quantum-mechanical coupling estimates. (*42, 43*)

### Marcus electron-transfer rate proxy

Relative electron-transfer rates were estimated using a Marcus-type expression that includes the driving force term:

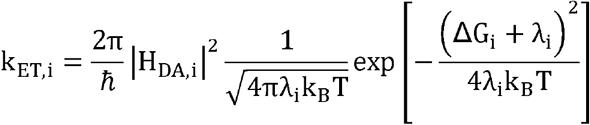

where \(H_{DA,i}\) is the donor–acceptor electronic-coupling proxy for hop \(i\), \(\lambda_i\) is the structure-derived reorganization-energy proxy, \(\Delta G_i\) is the driving force for the redox hop, \(k_B\) is the Boltzmann constant, and \(T\) is temperature.

Electronic coupling was modeled as

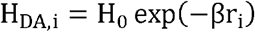

where r_i_ is the donor–acceptor distance, H_0_ is the coupling prefactor, and β is the distance-decay coefficient. Unless otherwise stated, calculations were performed using H_0_=0.010 eV, β = 1.1 \AA^−1^, ΔG = −0.08 eV, and T=300 K, k_B_T = 0.02585 eV. The reported rates should therefore be interpreted as relative structure-informed Marcus-type rate proxies rather than experimentally calibrated absolute electron-transfer rate constants.(*41*)

### Identification of electron-transfer bottlenecks

For each sequential redox hop, a bottleneck score was calculated:

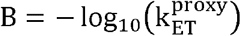

Higher bottleneck scores correspond to slower predicted electron transfer. To avoid artifacts arising from unrealistically long electron-transfer distances, hops exceeding 30 Å were excluded from bottleneck ranking analyses. This criterion excluded the SF4_C_301 → FES_O_301 transition (76.98 Å), which lies well beyond the distance range typically associated with efficient biological electron transfer. The hop exhibiting the largest bottleneck score was designated the dominant kinetic bottleneck.(*44*)

### Sensitivity analysis of the dominant bottleneck

To evaluate the robustness of the identified bottleneck, systematic perturbation analysis was performed on the H5 transition. Electronic coupling H_DA_ was varied from 0.1-fold to 100-fold of its observed value, whereas the reorganization-energy proxy λ was varied from 50% to 150% of baseline. For every parameter combination, the Marcus rate proxy was recalculated, generating a two-dimensional sensitivity landscape describing the dependence of electron-transfer efficiency on coupling and environmental reorganization.(*45*)

### Rescue simulations

To estimate the theoretical reversibility of the dominant bottleneck, rescue simulations were performed by increasing electronic coupling and decreasing reorganization energy. H_DA_ was increased by factors of 3, 10, 30, and 100, while (\lambda) was reduced by 9%, 19%, 30%, and 40%. For each perturbation,

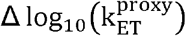

was calculated relative to the baseline H5 value. These analyses quantify the extent to which local structural modifications could theoretically alleviate the dominant electron-transfer bottleneck. (*18*)

### Open Quantum Transport Analysis

To investigate quantum transport behavior along the mitochondrial Complex I electron-transfer chain, we performed open quantum transport simulations using a Lindblad master-equation formalism. The electron-transfer pathway was represented as a site-based network consisting of FMN, Fe–S clusters, and the terminal quinone acceptor. Effective electronic couplings between neighboring redox centers were estimated from structure-derived distance-dependent coupling proxies.

Quantum-state evolution was described by

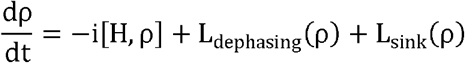

where ρ is the reduced density matrix, *H* is the effective site Hamiltonian, L_dephasing_ represents environmentally induced dephasing, and L_sink_ represents irreversible population transfer to the terminal acceptor state. Site-dependent dephasing rates γ were parameterized using local structural microenvironment descriptors derived from the protein structure.

The terminal quinone site was modeled as an irreversible sink, allowing transport efficiency to be quantified as the final sink population reached during numerical integration of the Lindblad equation. Although the underlying quantum dynamics were explicitly modeled through a Lindblad master-equation formalism, Hamiltonian couplings and environmental parameters were estimated from structure-derived proxy quantities rather than first-principles electronic-structure calculations. This framework enabled evaluation of how local structural constraints and bottleneck regions influence quantum transport efficiency along the Complex I redox pathway.

### Statistical analysis

All structural descriptors were converted into z-scores before comparison. Group comparisons were performed using two-sided non-parametric tests. Statistical significance was defined as (P<0.05). All analyses were conducted using Python 3.11 with NumPy, Pandas, SciPy, and Matplotlib. Figures were generated using publication-quality vector graphics and exported at 600 dpi resolution.

## Declarations

### Ethics approval and consent to participate

N/A

### Consent for publication

N/A

### Availability of data and materials

The analysis codes and datasets used and/or generated during the current study are available from Dr. Ji-Yong Sung upon reasonable request. Interested researchers may contact Dr. Sung via email at to obtain access to the relevant materials.

### AI Use Declaration

AI tools were used only for English grammar correction and language polishing.

### Authors’ contributions

Conceptualization & Investigation: JYS, JHC; Methodology: JYS; Data analysis: JYS; Writing-original draft: JYS; Writing-review & editing: JYS, JHC; Supervision: JYS, JHC; Project administration: JYS, JHC; Funding acquisition: JHC; Interpretation of the results; JYS, JHC. All authors have read and agreed to the published version of the manuscript.

## Acknowledgements

N/A

## Reference

1. G. Magro, V. Laterza, F. Tosto, Leigh Syndrome: A Comprehensive Review of the Disease and Present and Future Treatments. Biomedicines 13, (2025).

2. A. B. Bakare, E. J. Lesnefsky, S. Iyer, Leigh Syndrome: A Tale of Two Genomes. Frontiers in Physiology 12, (2021).

3. Z. Yin, A.-N. A. Agip, H. R. Bridges, J. Hirst, Structural insights into respiratory complex I deficiency and assembly from the mitochondrial disease-related ndufs4−/− mouse. The EMBO Journal 43, 225–249 (2024).

4. K. Fiedorczuk, L. A. Sazanov, Mammalian Mitochondrial Complex I Structure and Disease-Causing Mutations. Trends in Cell Biology 28, 835–867 (2018).

5. R. G. Efremov, L. A. Sazanov, The coupling mechanism of respiratory complex I — A structural and evolutionary perspective. Biochimica et Biophysica Acta (BBA) - Bioenergetics 1817, 1785–1795 (2012).

6. V. R. I. Kaila, Long-range proton-coupled electron transfer in biological energy conversion: towards mechanistic understanding of respiratory complex I. Journal of The Royal Society Interface 15, (2018).

7. C. C. Moser, J. M. Keske, K. Warncke, R. S. Farid, P. L. Dutton, Nature of biological electron transfer. Nature 355, 796–802 (1992).

8. H. R. Williamson, B. A. Dow, V. L. Davidson, Mechanisms for control of biological electron transfer reactions. Bioorganic Chemistry 57, 213–221 (2014).

9. M. L. Jones, I. V. Kurnikov, D. N. Beratan, The Nature of Tunneling Pathway and Average Packing Density Models for Protein-Mediated Electron Transfer. The Journal of Physical Chemistry A 106, 2002–2006 (2002).

10. M. Weik et al., Specific protein dynamics near the solvent glass transition assayed by radiation-induced structural changes. Protein Science 10, 1953–1961 (2008).

11. E. A. Dolan, R. B. Yelle, B. W. Beck, J. T. Fischer, T. Ichiye, Protein Control of Electron Transfer Rates via Polarization: Molecular Dynamics Studies of Rubredoxin. Biophysical Journal 86, 2030–2036 (2004).

12. D. R. Martin, D. V. Matyushov, Electron-transfer chain in respiratory complex I. Scientific Reports 7, (2017).

13. I. E. T. Iben et al., Glassy behavior of a protein. Physical Review Letters 62, 1916–1919 (1989).

14. H. Frauenfelder, S. G. Sligar, P. G. Wolynes, The Energy Landscapes and Motions of Proteins. Science 254, 1598–1603 (1991).

15. J. Zhu, K. R. Vinothkumar, J. Hirst, Structure of mammalian respiratory complex I. Nature 536, 354–358 (2016).

16. K. Rother, R. Preissner, A. Goede, C. Frömmel, Inhomogeneous molecular density: reference packing densities and distribution of cavities within proteins. Bioinformatics 19, 2112–2121 (2003).

17. F. Hoeser, P. Saura, C. Harter, V. R. I. Kaila, T. Friedrich, A leigh syndrome mutation perturbs long-range energy coupling in respiratory complex I. Chemical Science 16, 7374–7386 (2025).

18. T. Hayashi, A. A. Stuchebrukhov, Electron tunneling in respiratory complex I. Proceedings of the National Academy of Sciences 107, 19157–19162 (2010).

19. N. Lambert et al., Quantum biology. Nature Physics 9, 10–18 (2012).

20. J. Y. Sung, J. H. Cheong, Quantum biology: From mechanisms to medicine. Clinical and Translational Medicine 16, (2026).

21. J. Y. Sung, J. H. Cheong, Quantum medicine: A quantum–mechanical framework for redox biology, disease and precision medicine. Clinical and Translational Medicine 16, (2026).

22. M. Schubert Baldo, L. Vilarinho, Molecular basis of Leigh syndrome: a current look. Orphanet Journal of Rare Diseases 15, (2020).

23. L. A. Sazanov, Respiratory Complex I: Mechanistic and Structural Insights Provided by the Crystal Structure of the Hydrophilic Domain. Biochemistry 46, 2275–2288 (2007).

24. J. M. Berrisford, L. A. Sazanov, Structural Basis for the Mechanism of Respiratory Complex I. Journal of Biological Chemistry 284, 29773–29783 (2009).

25. E. Gnandt, K. Dörner, M. F. J. Strampraad, S. de Vries, T. Friedrich, The multitude of iron– sulfur clusters in respiratory complex I. Biochimica et Biophysica Acta (BBA) - Bioenergetics 1857, 1068–1072 (2016).

26. D. N. Beratan, J. N. Onuchic, J. R. Winkler, H. B. Gray, Electron-Tunneling Pathways in Oroteins. Science 258, 1740–1741 (1992).

27. H. B. Gray, J. R. Winkler, Electron Transfer in Proteins. Annual Review of Biochemistry 65, 537–561 (1996).

28. M. Karplus, D. Vitkup, D. Ringe, G. A. Petsko, Solvent mobility and the protein ‘glass’ transition. Nature Structural Biology 7, 34–38 (2000).

29. N. Kurochkina, G. Privalov, Heterogeneity of packing: Structural approach. Protein Science 7, 897–905 (2008).

30. J. C. Gaines, W. W. Smith, L. Regan, C. S. O’Hern, Random close packing in protein cores. Physical Review E 93, (2016).

31. J. Liang, K. A. Dill, Are Proteins Well-Packed? Biophysical Journal 81, 751–766 (2001).

32. A. Shabani et al., in Quantum Effects in Biology. (2014), chap. 2, pp. 14–52.

33. R. Dorner, J. Goold, L. Heaney, T. Farrow, V. Vedral, Effects of quantum coherence in metalloprotein electron transfer. Physical Review E 86, (2012).

34. R. Guo, S. Zong, M. Wu, J. Gu, M. Yang, Architecture of Human Mitochondrial Respiratory Megacomplex I2III2IV2. Cell 170, 1247–1257.e1212 (2017).

35. J. Hirst, Mitochondrial Complex I. Annual Review of Biochemistry 82, 551–575 (2013).

36. N. J. Lake, A. G. Compton, S. Rahman, D. R. Thorburn, Leigh syndrome: One disorder, more than 75 monogenic causes. Annals of Neurology 79, 190–203 (2015).

37. K. Fiedorczuk et al., Atomic structure of the entire mammalian mitochondrial complex I. Nature 538, 406–410 (2016).

38. K. L. Ngai, S. Capaccioli, N. Shinyashiki, The Protein “Glass” Transition and the Role of the Solvent. The Journal of Physical Chemistry B 112, 3826–3832 (2008).

39. A. Bondi van der Waals Volumes and Radii. The Journal of Physical Chemistry 68, 441–451 (2002).

40. J. Tsai, R. Taylor, C. Chothia, M. Gerstein, The packing density in proteins: standard radii and volumes 1 1Edited by J. M. Thornton. Journal of Molecular Biology 290, 253–266 (1999).

41. R. A. Marcus, On the Theory of Oxidation-Reduction Reactions Involving Electron Transfer. I. The Journal of Chemical Physics 24, 966–978 (1956).

42. H. B. Gray, J. R. Winkler, Electron tunneling through proteins. Quarterly Reviews of Biophysics 36, 341–372 (2004).

43. J. Winkler, Electron tunneling pathways in proteins. Current Opinion in Chemical Biology 4, 192–198 (2000).

44. C. C. Moser, S. E. Chobot, C. C. Page, P. L. Dutton, Distance metrics for heme protein electron tunneling. Biochimica et Biophysica Acta (BBA) - Bioenergetics 1777, 1032–1037 (2008).

45. J. Blumberger, Recent Advances in the Theory and Molecular Simulation of Biological Electron Transfer Reactions. Chemical Reviews 115, 11191–11238 (2015).

